# Widespread localisation of lncRNA to ribosomes: Distinguishing features and evidence for regulatory roles

**DOI:** 10.1101/013508

**Authors:** Juna Carlevaro-Fita, Anisa Rahim, Roderic Guigo, Leah A. Vardy, Rory Johnson

## Abstract

The function of long noncoding RNAs (lncRNAs) depends on their location within the cell. While most studies to date have concentrated on their nuclear roles in transcriptional regulation, evidence is mounting that lncRNA also have cytoplasmic roles. Here we comprehensively map the cytoplasmic and ribosomal lncRNA population in a human cell. Three-quarters (74%) of lncRNAs are detected in the cytoplasm, the majority of which (62%) preferentially cofractionate with polyribosomes. Ribosomal lncRNA are highly expressed across tissues, under purifying evolutionary selection, and have cytoplasmic-to-nuclear ratios comparable to mRNAs and consistent across cell types. LncRNAs may be classified into three groups by their ribosomal interaction: non-ribosomal cytoplasmic lncRNAs, and those associated with either heavy or light polysomes. A number of mRNA-like features destin lncRNA for light polysomes, including capping and 5’UTR length, but not cryptic open reading frames or polyadenylation. Surprisingly, exonic retroviral sequences antagonise recruitment. In contrast, it appears that lncRNAs are recruited to heavy polysomes through basepairing to mRNAs. Finally, we show that the translation machinery actively degrades lncRNA. We propose that light polysomal lncRNAs are translationally engaged, while heavy polysomal lncRNAs are recruited indirectly. These findings point to extensive and reciprocal regulatory interactions between lncRNA and the translation machinery.

## Introduction

The past decade has witnessed the discovery of a tens of thousands of long nonprotein coding RNAs (lncRNAs) in our genome, with profound implications for our understanding of molecular genetics, disease and evolution. Focus is now shifting to understanding the function to these molecules. We reason that such function is likely to be intimately linked to the location of lncRNA within the cell.

Following the first compelling discoveries of chromatin regulatory lncRNAs such as XIST (1)and HOTAIR (2), a paradigm was established for lncRNAs as nuclear-restricted, epigenetic regulatory molecules (3). However, it is not clear to what extent this is true for the >10,000 lncRNAs that remain uncharacterised (4-7). Indeed growing evidence points to lncRNA having diverse roles outside of the cell nucleus, including regulation of microRNA activity (8), protein sequestration(9), and mRNA translation(10).

Somewhat paradoxically, cytoplasmic lncRNA has recently been reported to interact with the ribosome. In footprinting experiments to map ribosome-bound transcripts genome-wide, the Weissman group identified a considerable number of lncRNAs directly engaged by the translation machinery (11), an observation subsequenttly corroborated in an independent study(12). These transcripts do not contain classical features of protein-coding sequence, and various analyses have argued that these lncRNAs are not productively translated in most cases (13,14). It is not yet clear whether ribosomal recruitment is a general property of all lncRNA in the cell. If not, it is of interest to understand what features distinguish ribosomal lncRNAs.

The biological significance of ribosomal lncRNA remains unclear. Two principle types of potential regulatory functions for ribosomal lncRNAs have been proposed: either sequence-specific regulation of mRNA translation or general regulation of ribosome function (15). LncRNA and mRNA arising from opposite genomic strands can form stable RNA-RNA hybrids that are localised in ribosomes (10). Through such “cis-antisense” interactions, lncRNA may specifically regulate stability and translation of their mRNA partner (10), although this has not yet been demonstrated at a genome-wide scale. The advent of ribosome footprinting technology has prompted the idea that lncRNA may non-specifically regulate translation through direct binding by ribosomes (15). Cryptic open reading frame (ORF) sequences within lncRNA may be recognised by the ribosome, directly resulting in translational repression or else enabling recruitment of regulatory proteins. Other more mundane scenarios are also possible: ribosomes might be a default destination of all polyadenylated mRNA-like transcripts, where they are recognised as non-coding and processed by one of various known quality surveillance pathways.

In the present study we take these studies further by comprehensively mapping the entire known cytoplasmic and ribosomal lncRNA population of a human cell line. We show that the majority of cytoplasmic lncRNAs are robustly and verifiably associated with ribosomes. We show evidence that lncRNAs can be divided into classes based on subcellular location and distinguished by a variety of features. These classes likely serve distinct regulatory roles in translation. Finally we show that the translation machinery serves as the endpoint of the lncRNA life-cycle. We conclude that, rather than being an exception, ribosomal recruitment is frequently the destination of cytoplasmic lncRNAs.

## Results

### Creating a high confidence lncRNA catalogue

Our aim was to map the distribution of lncRNAs in the cytoplasm and on the polysomes of human cells. A potential confounding factor in any analysis of ribosome-bound RNAs is the possibility of misannotated protein-coding transcripts (16). These represent a non-negligible fraction of lncRNA annotation, due to the technical challenges of correctly identifying protein coding sequences with high sensitivity, as well as biological factors: a number of annotated lncRNAs have subsequently been found to encode peptides, including small “micropeptides”, which were overlooked by conventional annotations (17,18).

We decided to implement the most stringent possible filtering to remove protein coding transcripts from our analysis, even at the expense of omitting some genuine non-coding transcripts. We first removed lncRNAs that could be unannotated extensions of protein-coding genes or pseudogenes. Remaining genes were filtered using a panel of methods for identifying protein coding sequence (Figure 1A and Materials and Methods). Altogether 9057 lncRNA transcripts (61.9%), 6763 genes (73.8%) were unanimously classified as non-coding - these we refer to as “filtered lncRNAs” (Figure 1A). The remaining genes of uncertain protein coding status are henceforth referred to as “potential protein coding RNAs” (4415 transcripts, 1878 genes). The complete sets of potential protein coding and filtered lncRNAs are available in Supplementary Table S1.

**Figure 1.**
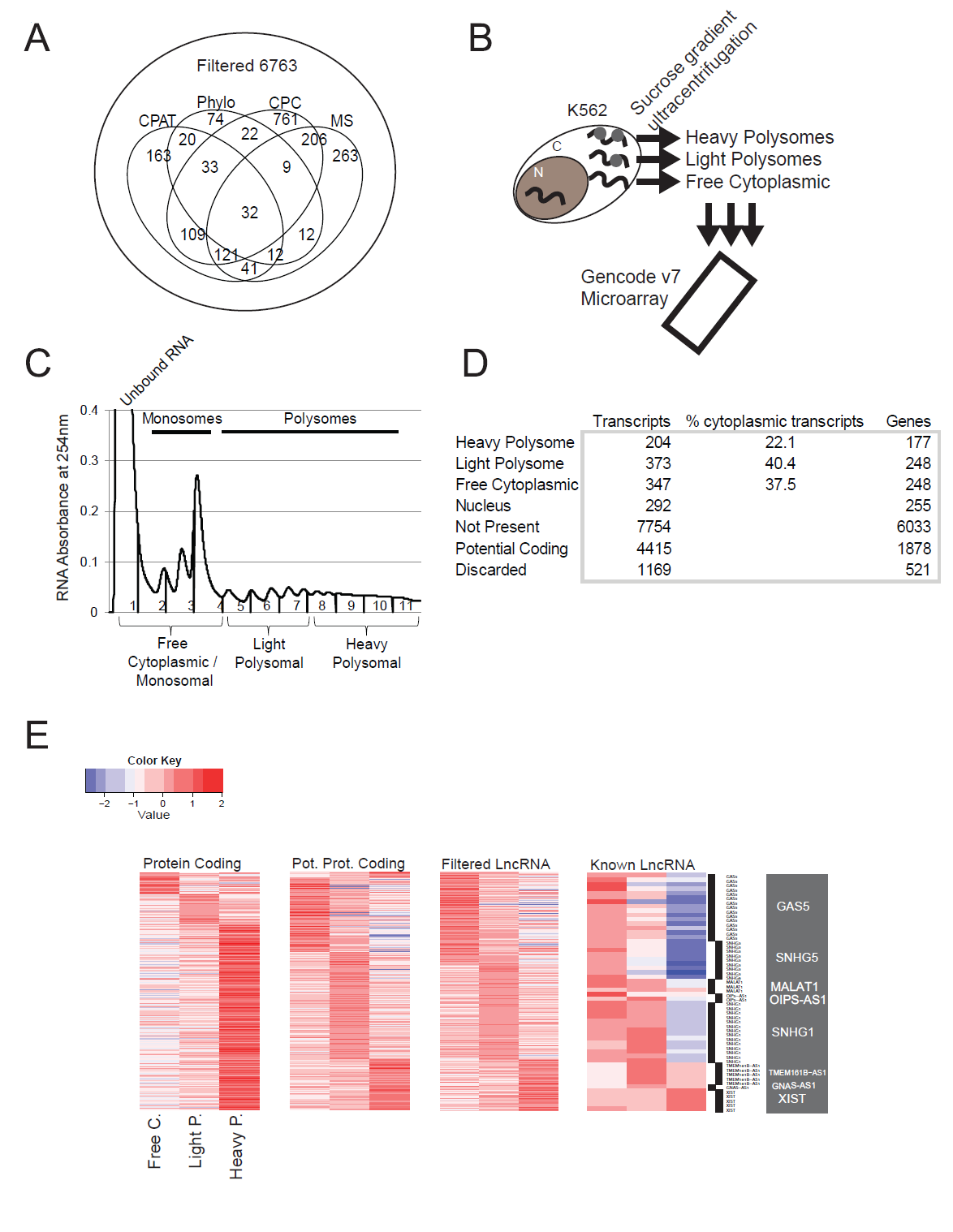
Legends Figure 1: Discovery and classification of ribosome-associated lncRNAs. (A) Numbers of Gencode v7 lncRNA genes filtered by protein-coding prediction methods. Genes (and all their constituent transcripts) having at least one transcript identified as protein-coding by at least one method were designated "Potential Protein Coding”. Remaining genes with no evidence for protein coding potential were defined as "Filtered LncRNAs”. (B) Outline of the subcellular mapping of K562 lncRNA by polysome profiling and microarray hybridisation. (C) Definition of the pooled fractions from sucrose ultracentrifugation used in this study. (D) Summary of the numbers of genes and transcripts classified in subcellular fractions. (E) Heatmaps show the relative microarray intensity measured for each RNA sample. The colour scale runs from blue (low detection) through white to red (high detection). “Protein coding” refers to the 2796 probes for protein-coding mRNAs included on the microarray, “Known lncRNAs” are those filtered transcripts that also belong to the lncRNAdb database(23).

### Mapping the cytoplasmic and ribosomal lncRNA population

We sought to create a comprehensive map of cytopasmic lncRNA localisation in a human cell. We chose as a model the K562 human myelogenous leukaemia cell line. Being an ENCODE Tier I cell, it has extensive transcriptomic, proteomic and epigenomic data publically available (19). We subjected cytoplasmic cellular extracts to polysome profiling, an ultracentrifugation method to identify ribosome-bound RNAs and distinguish transcripts bound to single or multiple ribosomes (Figure 1B) (20). Extracts were divided into three pools: “Heavy Polysomal”, corresponding to high molecular weight complexes cofractioning with >6 ribosomes; “Light Polysomal”, cofractioning with 2-6 ribosomes; and low molecular weight complexes corresponding to non-translated, cytoplasmic RNAs (Figure 1C). The latter contains free mRNAs found in the high peak in fraction 1, the 40 and 60S ribosomal subunits (fractions 2 and 3) and mRNAs that are bound by a single ribosome (fraction 4) - we define these as “Free Cytoplasmic” throughout the paper. It is important to note that although this fraction includes some RNAs bound by ribosomal subunits, or individual ribosomes, the majority of these are not considered to be efficiently translated (20).

Custom microarrays probing the entire Gencode v7 long noncoding RNA catalogue were used to analyse RNAs in the free cytoplasmic, light and heavy polysome fractions, in addition to total input RNA (see Materials and Methods)(5). Microarrays also contained probes targeting 2796 protein-coding genes. High positive correlation was observed between microarray RNA concentration measurements and RNA-sequencing of the same cells from ENCODE (Supplementary Figure S1)(19). Correlation between microarray results and RNAseq measurements of cytoplasmic RNA was higher than with either nuclear or whole-cell RNA from the same cells, attesting to the purity of these cytoplasmic extracts (Supplementary Table S2). Using stringent cutoffs we detected 10.6% of filtered lncRNA transcripts (962 transcripts, representing 665 or 9.8% of genes) and 52.8% of mRNAs (1476) in K562 cytoplasm (Figure 1D). An additional 292 transcripts (3.2%, representing 255 or 3.7% genes) were detected only in the nucleus. Altogether, 1254 filtered lncRNA transcripts (13.9%, representing 875 or 13.0% of genes) were detected.

We classified cytoplasmic lncRNAs according to their maximal ribosomal association, resulting in 347 (37.6% of cytoplasmic lncRNA transcripts) Free Cytoplasmic, 373 (40.4%) Light Polysomal, and 204 (22.1%) Heavy Polysomal transcripts (Figure 1D). Altogether, 62.5% of lncRNA transcripts detected in the cytoplasm have maximal detection in Light or Heavy Polysomal fractions. Two lines of evidence support this classification approach. First, 75% (959/1287) of protein-coding mRNAs are classified as Heavy Polysomal, consistent with their being actively translated and in accordance with previous studies (Figure 1E)(20,21). Second, protein abundance measurements show that Heavy Polysomal mRNAs are translated most efficiently (Supplementary Table S3) (22). In contrast, potential protein-coding transcripts had a similar global ribosome-association profile to filtered lncRNA, suggesting that they are not translated efficiently and underlining the stringency of our lncRNA filtering (Figure 1E). Ribosomal lncRNA are not apparently enriched for those that produce small peptides (Supplementary Table S4).

Cytoplasmic and ribosomal localisation has previously been reported for a number of lncRNA. To test the degree of agreement between these and our data, we examined the 297 lncRNA transcripts (from 60 genes) from the LncRNA Database (23) that are also present in the Gencode v7 annotation. SNHG5 (5) and Gas5 (9) were detected in the cytoplasm and classified as Free Cytoplasmic transcripts, consistent with previous reports. The snoRNA host Gas5 has previously been reported as associated with ribosomes (24). Although we classified this gene as Free Cytoplasmic based on its maximal detection, 11 out of 16 transcript isoforms of Gas5 were also clearly detected in Light and Heavy polysomal fractions although with lower microarray probe intensities. SNHG1 is another snoRNA host reported to be bound by ribosomes (25), which we classify in the Light Polysomal fraction. For other known lncRNAs, we map their subcellular location for the first time: GNAS-AS1 (Nespas) and MEM161B-AS1 are specifically associated with the Light Polysomal fraction.

### Independent evidence for ribosomal interaction of lncRNA

We next looked for additional evidence to support ribosomal interaction of lncRNA. During ultracentrifugation, it is possible that lncRNAs associated with non-ribosomal, high molecular weight complexes may co-sediment with polyribosomes and thus represent false positives. To investigate this, we repeated polysome profiling on cells treated with puromycin (puro), a drug that disrupts ribosomes, and profiled a set of candidate transcripts by volume-normalised RT-PCR (Figure 2A, B). Bona fide ribosome-bound transcripts are expected to relocalise to the free cytoplasmic fraction in response to puromycin. Eleven out of 16 (69%) ribosomal lncRNAs were validated in this way, similar to the 4/4 protein coding mRNAs tested. In contrast, 2/3 Free Cytoplasmic lncRNAs we examined showed minimal response to puro treatment. Thus in the majority of cases, cosedimentation reflects a physical interaction between lncRNA and ribosomes.

**Figure 2.**
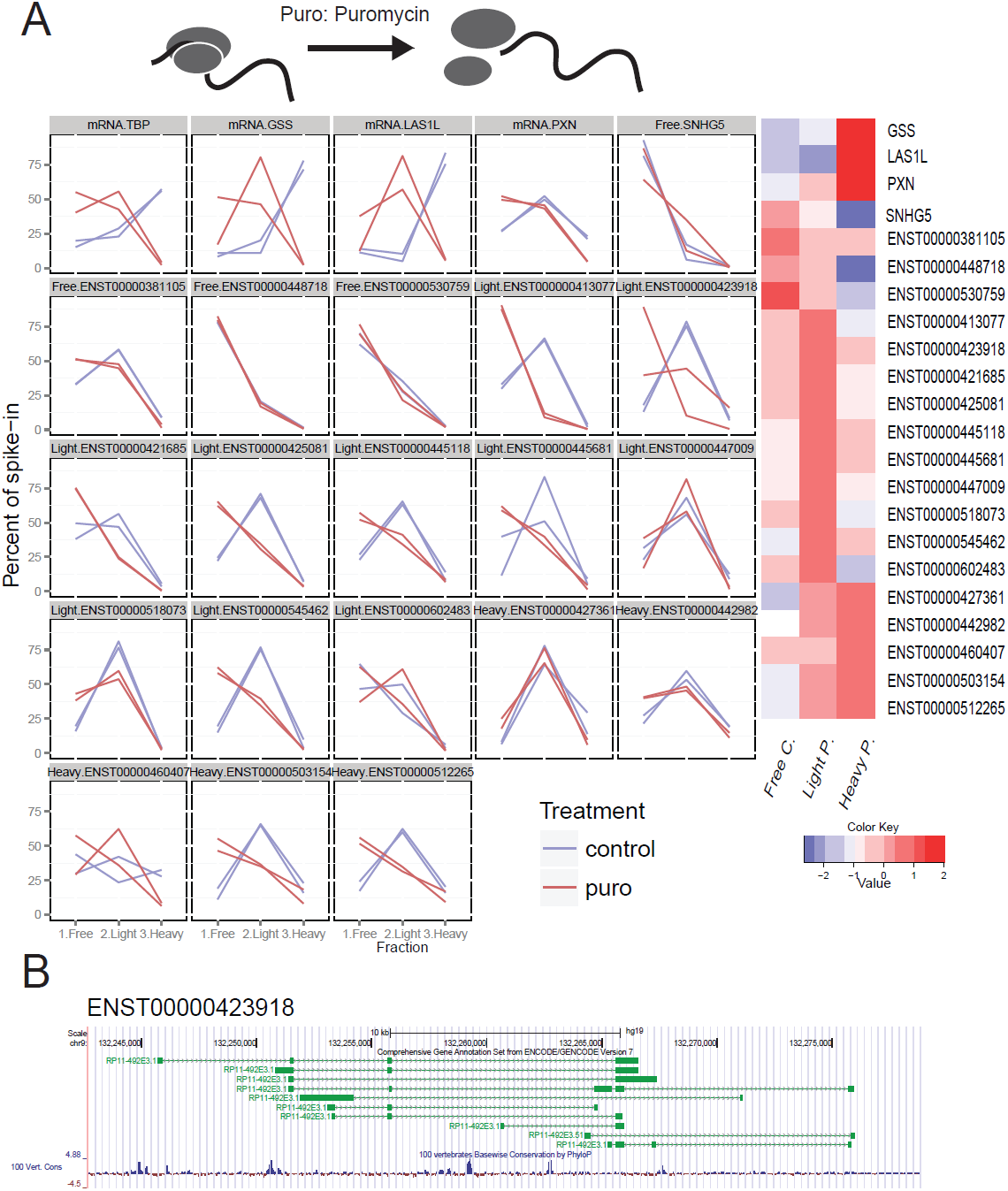
Validation of selected ribosome-associated lncRNA candidates. (A) We individually validated nine predicted ribosome-associated lncRNAs in independent ribosome profile experiments. In each case, two replicate experiments each were carried out with control K562 (red) and cells treated with puromycin (blue), for three distinct RNA fractions: (from left to right) free cytoplasmic, light polysomal, heavy polysomal. RNA levels are normalized to absolute levels of an RNA spiked into equal volumes of RNA sample. The first four panels represent protein coding mRNAs. Transcript IDs and classifications are shown above each panel. (B) Genomic map of ENST00000423918, a ribosome-associated transcript validated in this way.

We performed additional validation using fluorescence in situ hybridisation (FISH) to visualise the localisation of lncRNA at subcellular resolution. We tested three Light Polysomal lncRNAs (Figure 3). ENST0000504230 displays diffuse cytoplasmic localisation and exclusion from nucleoli. In addition to cytoplasmic localisation, the snoRNA precursor transcript ENST00000545440 (SNHG1) shows pronounced concentrations around the periphery of the nucleus, likely to be endoplasmic reticulum, and at three nuclear loci – possibly its site of transcription, given that the HeLa genome is predominantly triploid (26). Finally, ENST00000545462 (previously described as HEIH, a prognostic factor in hepatocellular carcinoma)(27), also has pronounced staining in the nuclear periphery, as well as within the nucleolus. Thus, both PCR and hybridisation methods support the interpretation from microarray data of ribosomal recruitment of lncRNA.

**Figure 3.**
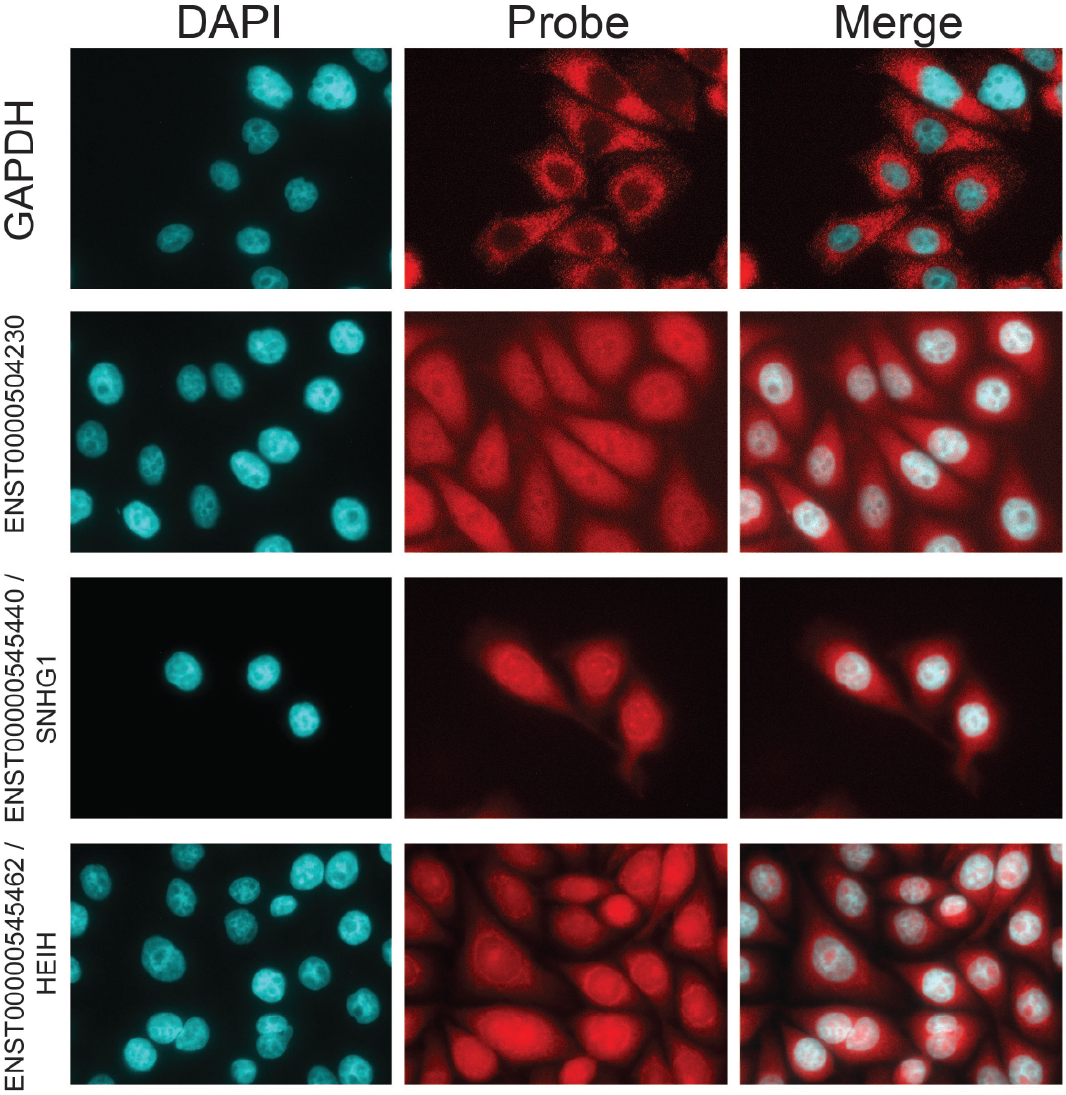
Fluorescence in situ hybridisation of ribosomal lncRNA in HeLa. Left Panel: DAPI staining of DNA; Middle: FISH probe; Right: merged. The actively translated housekeeping mRNA GAPDH was tested as a positive control for cytoplasmic localisation.

### Evidence for conserved function of ribosomal lncRNAs

Purifying evolutionary selection represents powerful evidence of functionality. A number of studies have shown that lncRNAs are under weak but non-neutral purifying evolutionary selection (5,28,29). We sought to test if this holds true for cytoplasmic lncRNAs, and in particular whether different classes of cytoplasmic lncRNA described above might have experienced different strengths of selection. We extracted PhastCons measures of exonic conservation and compared lncRNAs of distinct subcellular origins (Figure 4). Ancestral repeats were treated as neutrally-evolving DNA for comparison. As expected, protein coding exons have highly elevated conservation. Free Cytoplasmic, Light Polysomal and nuclear lncRNAs exhibit similar rates of non-neutral evolution. In addition, Heavy Polysomal lncRNAs contain a subset (∼10%) of transcripts with elevated conservation, second only to the potential protein coding transcripts, and higher than other expressed lncRNAs (P= 0.002, OR=2.40 Fisher test, testing the top 10% of Heavy Polysomal lncRNA vs other lncRNAs pooled). Thus, cytoplasmic lncRNAs experience purifying evolutionary selection consistent with conserved function.

**Figure 4.**
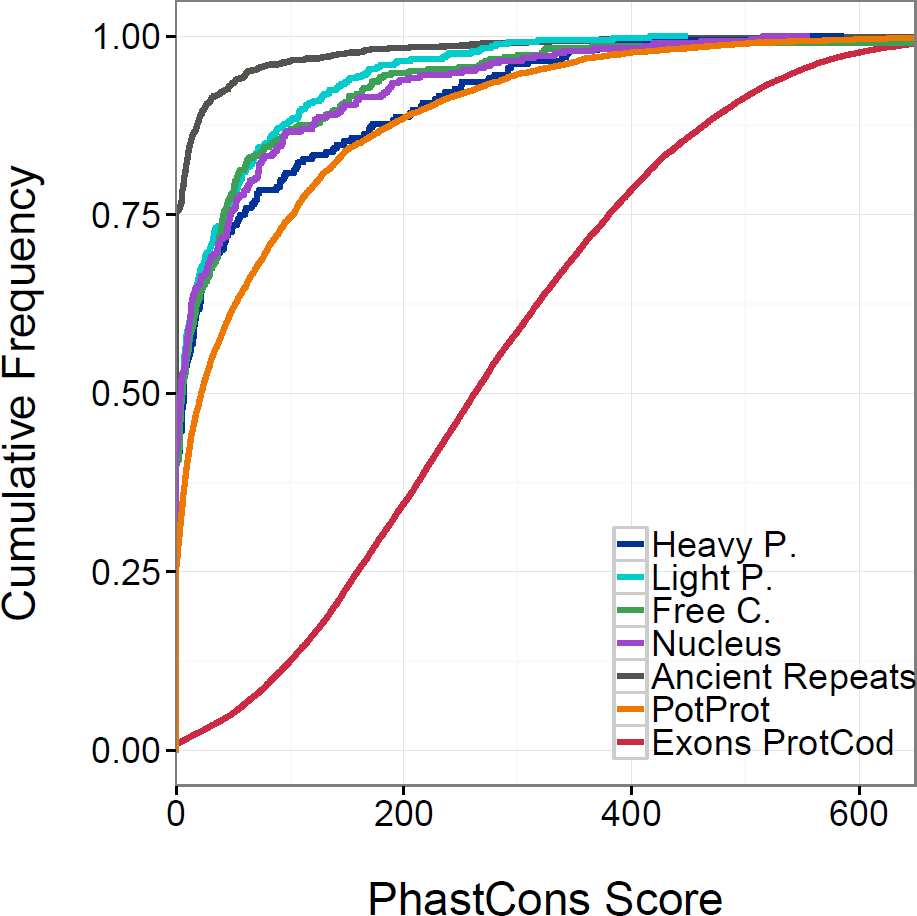
Ribosomal and cytoplasmic lncRNA are under purifying selection. Cumulative distribution of the mean PhastCons nucleotide-level conservation for the exons of the indicated transcript classes. PhastCons scores for ancestral repeats regions are also included to represent neutral evolutionary rates.

### Ribosomal lncRNA are highly expressed and consistently localised across cell types

We next investigated the organismal and subcellular expression patterns of lncRNA, in addition to their post-transcriptional processing. The steady state expression levels of cytoplasmic lncRNA is similar across cytoplasmic classes in independent K562 whole cell RNAseq, similar to that of mRNAs and well above nuclear-specific lncRNA (Figure 5A)(19). A similar trend is observed in human tissues: mean RPKM across Human Body Map tissues for all three cytoplasmic classifications exceed nuclear RNA (P = 5e-5 / 2e-5 / 0.0009 for Light Polysomal / Heavy Polysomal / Free Cytoplasmic vs nuclear, Wilcox test) (Supplementary Figure S2).

**Figure 5.**
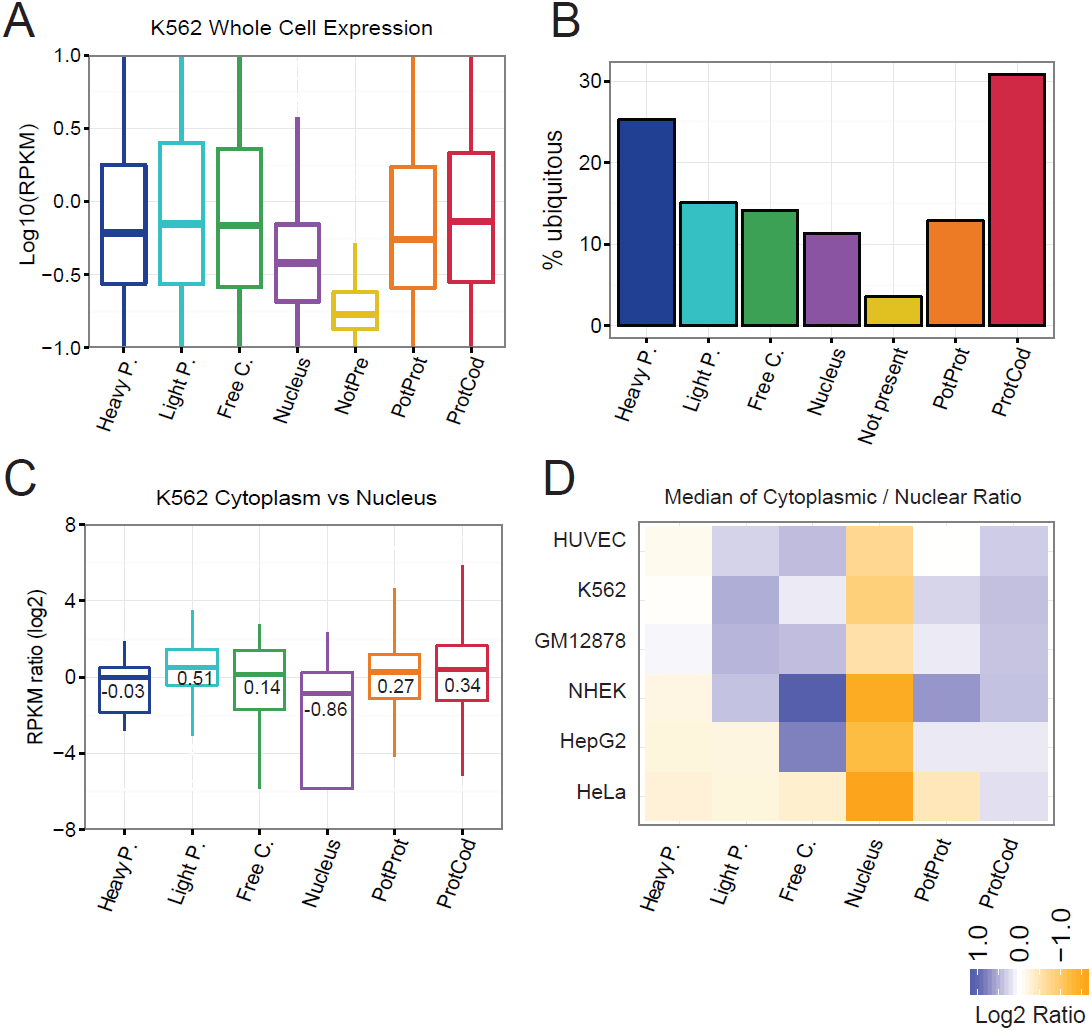
Intra-and Sub-cellular expression of ribosomal lncRNAs. (A) Expression in K562 whole cell by RNAseq. (B) Percent of transcripts having ubiquitous expression across human tissues defined by Human Body Map RNAseq. (B) Percent of ubiquitously expressed transcripts in each class. (C) Log2 cytoplasmic/nuclear RPKM ratios calculated from ENCODE RNAseq for indicated RNAs in K562 (whole cell, polyA+). For potential protein coding transcripts and mRNAs, data is only shown for detected transcripts. Numbers indicate median value. (D) Subcellular localisation of lncRNA amongst different cell lines. Colours reflect median cytoplasmic / nuclear RPKM values.

LncRNAs have been reported to be more tissue specific than mRNAs (4,5). Analysis of ubiquity, an inverse measure of tissue-specificity, of lncRNA in human tissues was consistent with this (Figure 5B). Despite this similarity in expression profiles, we find Heavy Polysomal lncRNAs to be significantly more ubiquitous in their tissue expression profiles compared to other lncRNA classes (P= 1.2e-4, Fisher Test Heavy Polysomal vs Nucleus), and essentially the same as mRNAs (P=0.129, Fisher exact test).

Subcellular localisation of lncRNA reported by polysome profiling is consistent with similar analysis using ENCODE RNAseq (19). Transcripts we report as ribosomal or free cytoplasmic have significantly elevated cytoplasmic-nuclear ratios (Figure 5C) (P=0.027 OR 2.4, 1.8e-07 OR 4.4, 0.005 OR 2.6 for Heavy Polysomal, Light Polysomal and Free Cytoplasmic vs Nuclear transcripts, Fisher exact test). Indeed, Light Polysomal lncRNA have median cytoplasmic specificity that exceeds protein coding mRNAs. Heavy Polysomal transcripts have a more nuclear distribution, suggesting that while some transcripts are ribosomally bound, other copies are present in the nucleus. We next asked whether the observed subcellular localisation of lncRNA in K562 is conserved across other cell types (Figure 5D). Similar analysis on RNAseq from other cell types showed Light Polysomal and Free Cytoplasmic transcripts tend to have high cytoplasmic-nuclear distributions, often exceeding that of mRNAs, while Heavy Polysomal has a more mixed distribution that nevertheless differs from nuclear-specific transcripts. Protein-binding profiles of lncRNA yields a consistent picture, with lncRNA tending to interact with proteins that localise to the same cellular compartment (Supplementary Figure S3). In summary, lncRNA subcellular localisation is consistent across cell types.

### mRNA-like 5’ regions distinguish ribosomally-bound lncRNAs

We next wished to identify factors that control the recruitment of lncRNA to ribosomes. The most obvious candidate feature is the ORF, especially given that lncRNAs contain abundant small ORF sequences that may be recognised by ribosomes. In protein, ORF length influences the number of ribosomes that can simultaneously bind, and hence the ribosomal fraction (compare mean sense ORF length for heavy and light polysome mRNA in Supplementary Figure S4)(12). However for lncRNA we could no evidence that ORFs determine ribosomal recruitment: neither their total ORF coverage, nor their number of ORFs, nor the length of their longest ORF is different from random sequence or correlates with ribosomal recruitment (Supplementary Figure S5). Nor apparently does gross gene structure or GC content, both clearly distinct between lncRNA and mRNA, appear to influence ribosomal recruitment (Supplementary Figure S6, S7).

We hypothesised that factors known to influence mRNA recognition by ribosomes may also apply to lncRNA. For mRNAs, a number of factors control the scanning and engagement by ribosomes, including 3’ polyadenylation, RNA structures within the 5’ UTR and 7-methylguanylate capping (30). To investigate whether polyadenylation influences ribosomal recruitment, we estimated the efficiency of polyadenylation of cytoplasmic and nuclear lncRNAs using ENCODE RNAseq on polyA+ and polyA-nuclear RNA. Although mRNA are more polyadenylated than lncRNA, we found no difference in polyadenylation efficiency between ribosomal and non-ribosomal lncRNAs (Supplementary Figure S8). We recently showed that splicing efficiency of lncRNAs is lower than mRNAs (31), but it does not distinguish ribosomal lncRNAs from other types (Supplementary Figure S9).

We next looked at the role of the 5’ end in ribosomal recruitment. Although lncRNAs do not have identifiable ORFs and hence 5’ UTRs, nevertheless they do contain abundant short “pseudo-ORFs”: random occurrences of in-frame start and stop codons. We defined the “pseudo-5’UTR” to be the region upstream of the first AUG trinucleotide of the lncRNA sequence. Although secondary structures in the 5’UTR have been shown to strongly influence translation of mRNAs (32), there is no overall difference in structural propensity between ribosomal and other lncRNAs (Supplementary Figure S10). However, the length of pseudo-5’UTRs does distinguish ribosomal from non-ribosomal lncRNA. Similar to protein-coding transcripts, Light Polysomal lncRNA have significantly longer 5’UTR regions than expected by chance (here estimated from the transcript’s reverse complement) (Figure 6A), while this effect is essentially absent for other cytoplasmic lncRNAs. Thus long 5’UTR-like regions would appear to contribute positively to ribosomal recognition of lncRNA.

**Figure 6.**
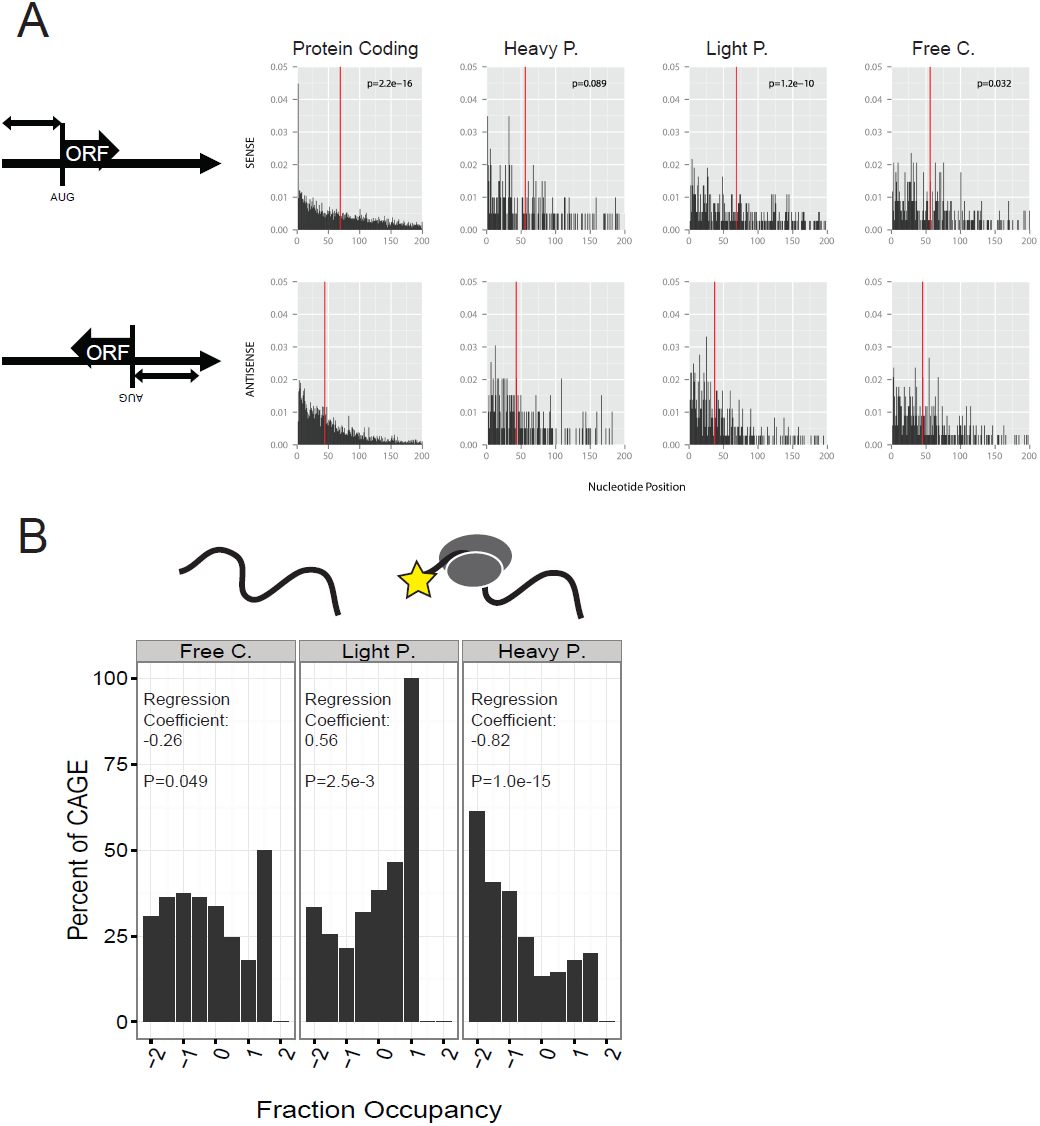
Ribosomal lncRNAs have mRNA-like 5’ ends. (A) The pseudo 5’ UTR was defined to be the distance from the start to the first AUG trinucleotide (top row). As a control, we calculated the same measure on the antisense strand (bottom row). Shown is the distribution of these lengths for each set of transcripts - protein coding mRNA (left), followed by cytoplasmic lncRNA classes. The red line indicates the mean value. P-values are for comparison of sense and antisense distributions using the Kolmogorov-Smirnov test. (B) Capping efficiency positively correlates with Light Polysome localisation of lncRNA. We defined every transcript to be capped if it has a K562 cytoplasmic polyA+ CAGE tag within 100bp upstream or downstream of its transcription start site. LncRNA were binned according to their relative enrichment in each of the three cytoplasmic fractions (x axis). In each bin, the percent of capped transcripts is shown in the y axis. Logistic regression was used to assess the relationship between these variables.

Recognition of the 5’ methyl-guanosine cap is required for mRNA scanning by the 40S ribosomal subunit. Using CAGE (cap analysis of gene expression) data (19), we examined the relationship between the ribosomal recruitment of lncRNA and capping using logistic regression. As shown in Figure 6B, there is a strong positive relationship between capping and recruitment to the Light Polysomal Fraction. In contrast, this relationship is negative for Free Cytoplasmic and Heavy Polysomal recruitment. This data suggests that capping of lncRNA is a driver of ribosomal recruitment, at least to the light polysomal fraction.

### Endogenous retroviral fragments are negatively correlated with ribosomal recruitment

There is growing evidence that transposable elements (TEs) contribute functional sequence to lncRNA (33,34). Taking all TE classes together, we observed an excess of TE-derived sequence within Free Cytoplasmic lncRNAs (P=4e-14, compared to remaining detected filtered lncRNAs, Wilcoxon test) (Figure 7A). Potentially protein coding transcripts are significantly depleted for TEs (P=2e-16, compared to all detected, filtered lncRNAs, Wilcoxon test). Given that protein coding transcripts are strongly depleted for TE insertions (35), this latter observation supports the idea that a subset of potential protein coding transcripts do indeed encode functional protein.

**Figure 7.**
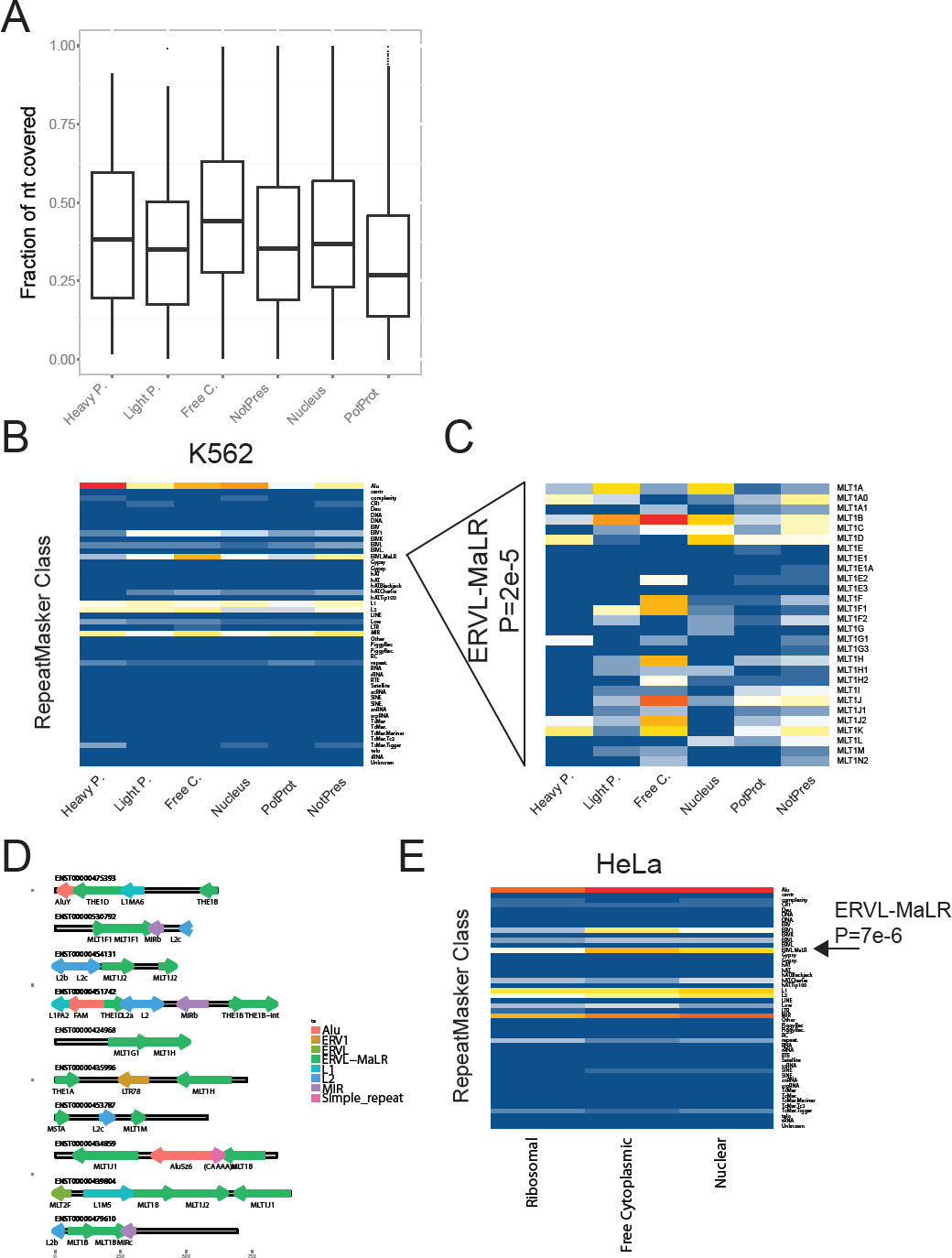
Transposable element composition of lncRNAs. (A) The fraction of each transcript covered by annotated repeat sequence from RepeatMasker. (B) The heatmap shows the normalised frequency of insertion for RepeatMasker-defined classes, ie the number of insertions per class divided by the length of each transcript. (C) As in (B) but showing data only for MLT-type repeats. (D) The repeat composition of a selection of Free Cytoplasmic, MLT-containing lncRNAs. The direction of the arrows indicates the annotated strand of the repeat with respect to the lncRNA. The colours represent the repeat class. (E) As in (B), except showing data for HeLa derived from ribosome footprinting experiments(36). For practical reasons, the lncRNAs are divided into three classes, see Materials and Methods for more details.

We were curious whether there exist TEs whose presence correlates with the subcellular localisation of their host transcript (Figure 7B) (Materials and Methods). Thus we systematically tested the relationship between subcellular localisation and TE class. We observed a relationship between the presence of Alu and transcript expression in K562: Alu are enriched amongst detected compared to undetected filtered lncRNAs (P=6e-7, Hypergeometric test), as recently described for human tissues (34). TcMar.Tigger, although rare, show evidence for preferential enrichment in polyribosomal lncRNAs (P=9e-4, Hypergeometric test). However the most obvious case is for the class of ERVL-MaLR, which are approximately two-fold enriched in free cytoplasmic lncRNAs compared to other expressed lncRNAs (Figure 7B). Closer inspection revealed that this effect is not due to a single repeat type, but rather to around a dozen subclasses of MST, MLT and THE endogenous retroelements (Figure 7C). We found no significant difference in the length of ERVL-MaLR insertions between lncRNA classes (Supplementary Figure S11). Rather it is the relative proportion of transcripts carrying an insertion that differs between groups. A selection of ERVL-MaLR containing lncRNAs are shown in Figure 7D.

Enrichment of ERVL-MaLR class elements in Free Cytoplasmic lncRNAs appears to be independent of cell type: using ribosome footprinting data from HeLa(36) we observe that ERVL-MaLR class TEs are specifically depleted from ribosome-bound lncRNAs (Figure 7E). Together these data suggest that endogenous retrovirus fragments may influence lncRNA trafficking in the cell.

### Evidence for cis-antisense lncRNA-mRNA pairing in ribosomes

Several reports exist describing hybridisation of lncRNA to mRNA through complementary sequences, resulting in trafficking of the former to ribosomes. Antisense complementarity between lncRNA and mRNA could take one of two forms: more conventionally, the two transcripts may originate from opposite strands of the same genomic locus, thus sharing complementary sequence regions (here “exonic antisense”, also referred to as “cis-antisense”) (Figure 8A) (10). More recently, it was shown that lincRNA-P21 contains regions of complementarity to mRNAs, through which they hybridise and consequently localise together in the ribosome (37). Importantly, the genes for lincRNA-P21 and its targets are located in distinct genomic loci – these we define here as “trans-antisense” pairs.

**Figure 8.**
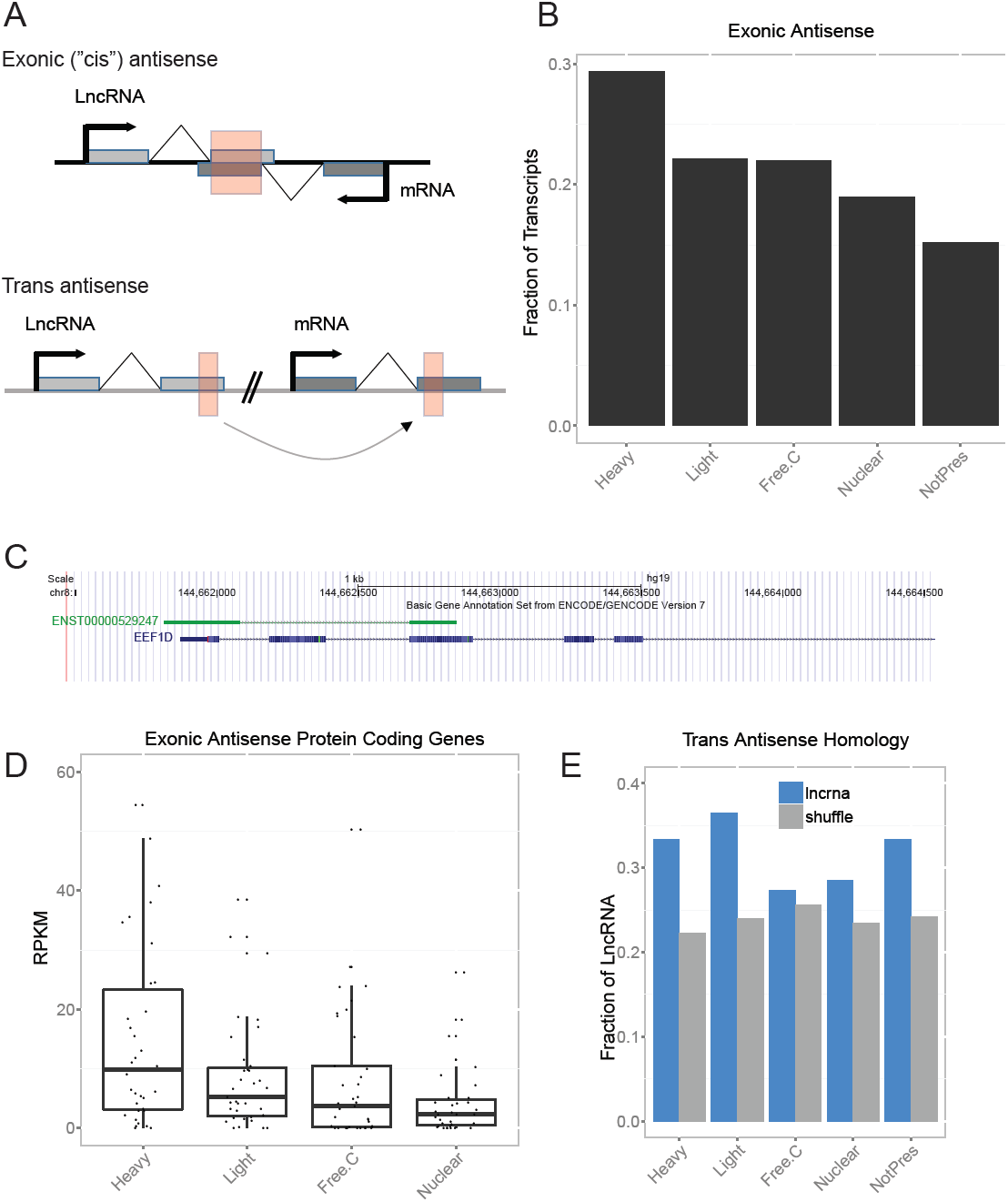
Cis- and Trans-antisense lncRNA-mRNA pairs and ribosomal recruitment. (A) Cartoon illustrating the definition of cis- and trans-antisense lncRNA-mRNA pairs. Red boxes indicate regions of opposite-strand homology. (B) The percent of each subcellular lncRNA class defined as exonic-antisense (cis-antisense) to a protein coding gene. (C) Example of a cis-antisense lncRNA-mRNA pair. ENST00000529247 (forward strand) is a heavy polysomal lncRNA transcribed antisense to the EEF1D gene (reverse strand), encoding a subunit of the translation elongation factor 1 complex. (D) Whole cell K562 polyA+ steady state levels of mRNAs that are antisense to lncRNA in the indicated subcellular classes. (E) The percent of lncRNA (blue bars) or size-matched random genomic fragments, having an antisense trans homology match to at least one mRNA.

We investigated whether either type of antisense may contribute to the observed recruitment of lncRNA to the ribosomes. We first hypothesised that exonic antisense lncRNAs would be more frequently localised in heavy polysomes, due to hybridisation to their corresponding (actively translated) mRNA. We classified all lncRNA by their genomic organisation with respect to protein-coding genes (5): intergenic (not overlapping), exonic antisense, intronic antisense, or intronic same sense. Consistent with our hypothesis, lncRNAs identified in heavy polysomes are significantly enriched for exonic antisense transcripts compared to those in other cellular compartments (P=4.2e-5, Fisher exact test) (Figure 8B). This finding is consistent with lncRNA / mRNA hybrids existing in human ribosomes. An example of such a cis-antisense pair is shown in Figure 8C. If this is the case, we would expect mRNAs bound by antisense heavy polysomal lncRNA to be more highly expressed than others. Examining RNAseq expression data we find this to be the case: mRNAs antisense to heavy polysomal lncRNA are significantly more highly expressed than mRNAs antisense to other lncRNA classes (P=7e-4, Wilcoxon test)(Figure 8D). In the course of this analysis, we also made the incidental observation that intronic same sense lncRNAs tend to be nuclear-specific (Supplementary Figure S12).

Trans-antisense hybridisation is another potential means by which lncRNA could interact with mRNA and be recruited to ribosomes. In this model, the transcripts share homology on opposite strands, but are not transcribed from the same genomic locus (Figure 8A). Using a BLAST approach, we compiled all sense-antisense homology relationships between intergenic lncRNA and mRNA (Figure 8E). As a control, we performed the same operation with size-matched, randomised genomic regions instead of lncRNA. This analysis resulted in two observations: first, lncRNA as a whole are more likely to have trans-antisense homology to mRNA compared to random genomic sequence (P=1e-14, Fisher test, comparing all lncRNA to all shuffled); second, this tendency was observed with statistical significance in ribosomal lncRNA (P=0.002, Fisher test for heavy and light lncRNA combined) but not in free cytoplasmic and nuclear lncRNA. These findings were consistent across a range of different BLAST settings. This data point to possible trans-antisense lncRNA-mRNA hybridisation as a general regulatory mechanism of ribosomal lncRNA.

### Degradation of lncRNA by the ribosome

We were next curious whether recruitment to ribosomes had any effect on lncRNA. It was proposed by Chew et al(38) that lncRNA at ribosomes are subject to nonsense mediated decay (NMD), and indeed one report does exist of ribosome-dependent degradation of a snoRNA host gene (35). We tested whether blocking translation has any outcome on the stability of ribosomal lncRNA identified here. Using the same candidate genes tested previously, we tested whether interfering with ribosomal function through drug-induced stalling (cyclohexamide) influenced lncRNA stability (Figure 9). In a number of cases we observed elongation-dependent degradation of lncRNA, often decreasing lncRNA amounts by several fold over 6 hours. This effect was highly heterogeneous, with other transcripts unaffected by cyclohexamide treatment. Thus, ribosomal recruitment leads to degradation of many lncRNAs.

**Figure 9.**
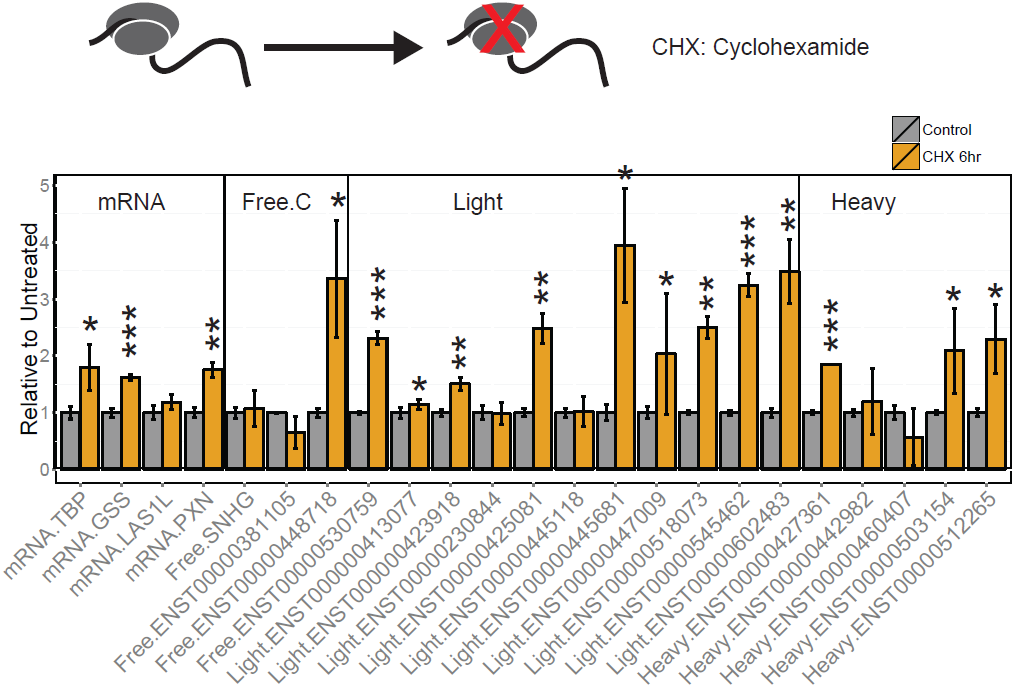
Changes in lncRNA stability in response to ribosome stalling. Bars represent mean detection in cells treated with cyclohexamide (CHX) for 6 hours and control cells (0h). Experiments were performed with three biological replicates. Bars show mean and standard deviation. Statistical significance was calculated by one-sided t-test (* P<0.05, ** P<0.01, *** P<0.001)

## Discussion

In order to gain clues as to lncRNA function at a global level, we have comprehensively mapped the ribosomal and cytoplasmic lncRNA populations of a human cell. The very substantial populations of lncRNA we discover in these fractions is at odds with existing paradigms of lncRNA function as principally nuclear molecules. We must now consider the possibility that lncRNA play more diverse roles outside the nucleus, including translational control, cellular metabolism or signal transduction.

One key challenge in the study of cytoplasmic lncRNAs is to rule out the possibility that they encode a cryptic, unannotated protein product. This question has been discussed in excellent reviews elsewhere (15), and has not yet been satisfactorily resolved. Indeed, it is likely that an extensive “grey zone” of transcripts with weak protein coding potential exists (and indeed may form the substrate for novel protein evolution (39)). It is also plausible that some or many transcripts do exist that function both as protein-coding and noncoding transcripts, although apart from the archetypal SRA1 (40), few concrete examples have so far been presented (41,42). In this study we took great pains to filter any transcripts with even minimal probability of encoding protein, in the process collecting many weakly coding (“potential protein coding”) transcripts that may be of rich scientific interest in future. We describe a set of 1867 annotated lncRNAs that have varying degrees of evidence for encoding protein. These transcripts have intriguing characteristics intermediate between coding and non-coding RNA: they are under higher evolutionary selection than lncRNA, are depleted for transposable elements, and tend to be cytoplasmically enriched – similar to mRNAs. In contrast, they have ribosomal association profiles and expression ubiquity similar to lncRNA. Finally, their gene structures and expression levels are intermediate between coding and noncoding sequences. It will be fascinating in future to study whether these transcripts represent an intermediate timepoint in the evolution of either new proteins from non-coding sequences, or the evolution of non-coding RNAs from formerly coding transcripts.

We cannot rule out the possibility that some of our filtered lncRNA produce a peptide product. Recent studies have revealed the potential for large volumes of unrecognised protein coding capacity in mammalian genomes, either as small peptides (43) or non-canonical ORF translation (44,45). However, even if these lncRNAs give rise to peptides, it does not necessarily follow that all are functional: occasional nonsense translation of ribosomally-localised lncRNA may occur with no functional consequences - “translational noise”. Future intensive mass spectrometry studies of short peptides, such as that carried out by Slavoff et al (18), will hopefully allow us to further improve lncRNA annotations.

We present several lines of evidence that ribosomal lncRNA are a large functional gene class that genuinely interacts with the translation machinery: (1) ribosomal lncRNA are puromycin sensitive; (2) fluorescence in situ hybridisation indicates their cytoplasmic localisation; (3) they have elevated cytoplasmic-nuclear ratios by independent ENCODE RNAseq data across diverse cell types; (4) their sequence is under similar or even elevated evolutionary selection compared to nuclear lncRNAs. Thus these transcripts are functional and appear to have a regulated and consistent subcellular localisation.

Polysome profiling appears to distinguish lncRNAs with distinct properties. We have attempted to rather crudely classify transcripts according to their fraction of maximum detection, but most transcripts are detected at varying concentrations in all fractions. Nevertheless, through this classification we have managed to discover features that distinguish lncRNA and laid a foundation for predicting lncRNA localisation *de novo*. Similarly, a recent study discovered an RNA motif that predicts and appears to confer cytoplasmic localisation (46). We find that lncRNAs localised in the Light Polysomal fraction tend to have mRNA-like 5’ features, more specifically a non-randomly long pseudo-5’ UTR length and the presence of a cap structure. This is consistent with the importance of 5’ recognition in the initiation of translation. Other mRNA-like features such as polyadenylation, GC content or open reading frames do not appear to affect ribosomal interaction at a global level. In contrast, repetitive sequence features, and particularly human endogenous retrovirus fragments, are negatively associated with ribosomal recruitment. This is perhaps to be expected, given that mRNAs tend to be depleted of such repeats, in contrast to lncRNA (35). The mechanism by which hERV prevent lncRNA from ribosomal recruitment remains to be ascertained, although we proposed recently that such fragments may interact with protein complexes that could antagonise ribosomal binding (33). In summary, these findings represent a starting point for discovering features that distinguish lncRNA classes and may eventually lead to useful models for predicting such classes.

Light Polysomal and Heavy Polysomal lncRNA appear to represent functionally distinct classes of lncRNA. It is generally considered that heavy polysomes tend to be actively translating, while light polysomes represent more weakly translated messengers, and this is supported by our mRNA data. We interpret the lncRNAs in the Light Polysomal fraction to be engaged by two or a few ribosomes that are not in the process of translating an mRNA. The above features of 5’ processing distinguish Light Polysomal transcripts clearly from Free Cytoplasmic transcripts, but not Heavy Polysomal. The latter tend to be more nuclear and more strongly evolutionarily conserved. Some clues to the origin of these differences may be gleaned from the observation that cis-antisense transcripts are enriched in the heavy fraction. Cis-antisense transcripts have been studied for a number of years, and cases have been described where the antisense lncRNA hybridises with its sense mRNA and accompanies it to the ribosome (10). Thus we might posit that lncRNA in heavy polysomes are involved in active translational processes and include transcripts that exist as hybrids with their sense mRNA partner. Such recognition is sequence specific, and we may guess that this localisation occurs indirectly: the lncRNA is recruited through its binding to a translated mRNA, and not directly engaged by ribosomes. In contrast, given their mRNA-like 5’ features, we propose that Light Polysomal lncRNA include cases that are directly engaged by ribosomes, resulting in non-specific translational repression and/or lncRNA degradation(15). This model is outlined in Figure 10.

**Figure 10.**
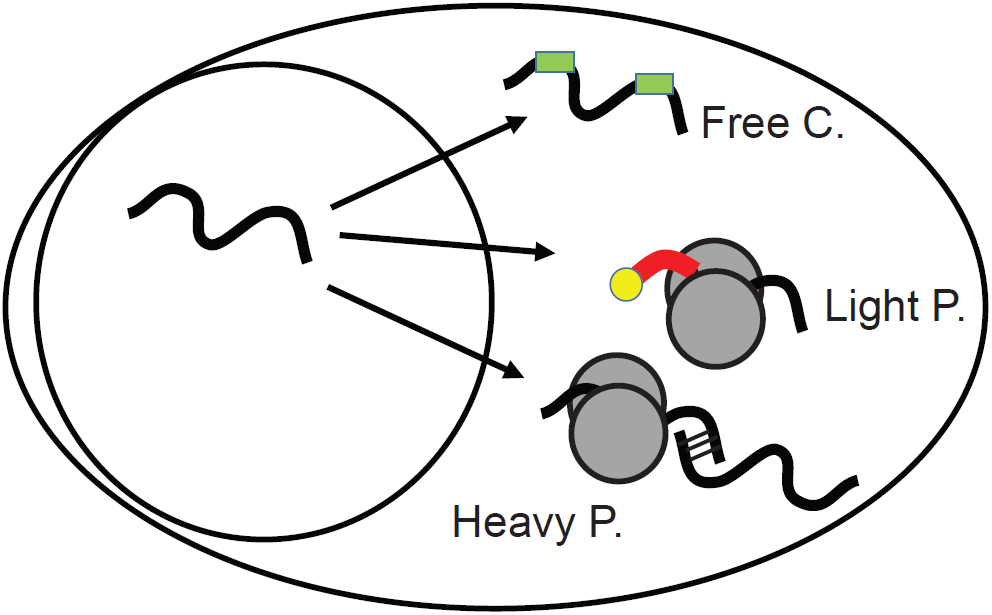
A model of lncRNA targeting within the cytoplasm.

Although it is tempting to propose that ribosomal lncRNA regulate protein translation, we must also seriously consider an alternative possibility: that the ribosome represents the default endpoint of the lncRNA lifecycle, and it is rather the non-ribosomal cytoplasmic transcripts that are exeptional. Indeed, it is perhaps not surprising that these mRNA-like transcripts - capped, polyadenylated and 100-10,000 nt long – should be recognised by the cell and trafficked accordingly. We here show evidence that, at least for a subset of transcripts, the result of ribosomal recruitment is degradation. That is, the translation machinery is also responsible for lncRNA clearance, and that the regulatory relationship between lncRNA and the translational machinery is reciprocal.

## Materials and Methods

### Polysome fractionation

For polysome fractionations, 20 million K562 cells were incubated with 100 ug/mL of cycloheximide (Sigma, Cat C4859) for 10 min. Cell pellets were resuspended in 200ul RSB buffer (20 mM Tris-HCl, pH 7.4, 20 mM NaCl, 30 mM MgCl_2_, 200ug/mL cycloheximide, 0.2mg/mL heparin (Sigma, Cat No. H4787), 1000 unit/mL RNasin), then lysed with an equal volume of Lysis Buffer (1X RSB, 1 % Triton X-100, 2% Tween-20, 200ug/ul heparin) with or without 1% Na deoxycholate. Following incubation on ice for 10 min, extracts were centrifuged at 13,000 x g for 3 min to remove the nuclei. Supernatants were further centrifuged at 13,000 x g for 8 min at 4°C. Equal OD units were loaded onto 10% to 50% linear sucrose gradients (prepared in 10 mM Tris-HCl pH 7.4, 75 mM KCl and 1.5mM MgCl_2_), and centrifuged at 36,000 rpm for 90 min at 8° C in a SW41 rotor (Beckman Coulter). Twelve fractions were collected from the top of the gradient using a piston gradient fractionator (BioComp Instruments). A UV-M II monitor (BIORAD) was used to measure the absorbance at 254 nm. 110ul of 10% SDS and 12 uL of proteinase K (10 mg/mL Invitrogen) was added to each 1ml fraction and incubated for 30 min at 42°C. Fractions 1-5, 6-8 and 9-11 were pooled corresponding to groups Free Cytoplasmic (Free / Monosomal), LP (Light Polysome) and HP (Heavy Polysome), respectively. For puromycin-treated samples, cells were incubated in 100ug/ml puromycin for 15 minutes prior to processing and puromycin was used in place of cyclohexamide in all the buffers.

Unfractionated cytoplasmic RNA and pooled polysomal RNAs were purified using phenol chloroform isoamyl extraction followed by LiCl precipitation to remove the heparin. The integrity of the samples was monitored by Bioanalyzer. For qRT-PCR analysis equal volumes of RNA were used to synthesise cDNA using the Superscript III Reverse Transcriptase (Invitrogen) according to manufacturer’s instructions. Two bacterial spike-in RNAs, Dap and Thr were added before RNA purification to equal volumes of each polysomal RNA pool. Gene specific primers were used with SYBR Green for qRT-PCR on an ABI PRISM 7900 Sequence Detection Systems. Candidate CT values were normalized to the spike in controls Dap and Thr that were present at equal concentrations per pool. Relative RNA levels are presented as a percentage of the RNA present in each pool with 100% RNA calculated as the sum of the FM, LP and HP pools.

### Microarray Design

This study was carried out using Agilent custom gene expression microarrays, in the 8x60k format with 60mer probes. Probes were designed using eArray software with standard settings: Base composition methodology / 60mer / 4 probes per target / sense probes / best probe methodology / 3' bias. Probes were designed for 14700 transcripts from the entire Gencode v7 lncRNA catalogue, in addition to 26 known lncRNAs from www.lncrnadb.org (23) and 90 randomly-selected protein-coding housekeeping genes. The array was then filled with probes targeting 2796 randomly-selected protein-coding gene probes. Microarray design details are available from the Gencode website (https://www.gencodegenes.org/lncrna_microarray.html).

### Microarray Hybridization and Probe Quantification

100 ng of total RNA was labeled using Low Input Quick Amp Labeling kit (Agilent 5190-2305) following manufacturer instructions. mRNA was reverse transcribed in the presence of T7-oligo-dT primer to produce cDNA. cDNA was then in vitro transcribed with T7 RNA polymerase in the presence of Cy3-CTP to produce labeled cRNA. The labeled cRNA was hybridized to the Agilent SurePrint G3 gene expression 8x60K microarray according to the manufacturer's protocol. The arrays were washed, and scanned on an Agilent G2565CA microarray scanner at 100% PMT and 3um resolution. Intensity data was extracted using the Feature Extraction software (Agilent).

Raw data was taken from the Feature Extraction output files and was corrected for background noise using the normexp method(47). To assure comparability across samples we used quantile normalization(48). All statistical analyses were performed with the Bioconductor project (http://www.bioconductor.org/) in the R statistical environment (http://cran.r-project.org/) (49).

### Preparation of filtered lncRNA gene catalogues

We first filtered the former set to remove any transcripts that potentially result from misannotated extensions or isoforms of protein-coding genes or pseudogenes. Any gene was discarded that has at least one transcript fulfilling one of the following conditions: overlapping on the same strand a Gencode v18 annotated pseudogene, overlapping on the same strand an exon of a protein-coding mRNA, or lying within 5 kb and on the same strand as an Gencode v18 protein-coding transcript or pseudogene (1169 transcripts, 521 genes). This resulted in a dataset of 13,472 lncRNA transcripts (8641 genes). Next, genes having at least one transcript predicted as protein coding by at least one method, were classified as “potential protein coding RNAs” (4415 transcripts, 1878 genes), while the remainder were classified as “filtered lncRNAs”. The four filtering methods used were: 1) PhyloCSF, a comparative genomics method based on phylogenetic conservation across species (50). The analysis was performed using 29 mammalian nucleotide sequence alignments and assessing the three sense frames. The alignment of each transcript was extracted from stitch gene blocks given a set of exons from Galaxy(51). Transcripts with score >95 were classified as potential protein coding, following the work of Sun et al (52) . 2) Coding Potential Assessment Tool (CPAT)(53), using the score threshold of 0.364 described by the authors. 3) Coding Potential Calculator (CPC), a support vector machine-based classifier based on six biological sequence features, using a cutoff of 1 (54). 4) Peptides: We used experimental mass spectrometry tag mappings from Pinstripe to identify any transcripts producing peptides (55). Any transcript having an exonic, same strand tag mapping were designated as “potential protein coding”. Collectively, sequence filters reduced the pool of analyzed transcripts to 9057 transcripts (6763 genes). The full table of classification data for all Gencode v7 lncRNA is available in Supplementary Table S1.

### Microarray probe filtering

LncRNA transcripts were considered to be present in a sample when at least three out of four microarray probes were reliable and not absent. The expression intensity value for “present” transcripts was computed as the median of its present probes. Protein coding genes were considered “present” if at least one probe was reliable and not absent, and the intensity value was that of one of the present probes, chosen randomly.

Variances in probe intensity values within probesets were significantly different when comparing all probsets from present transcripts in a sample (Levene’s test). To avoid non representative intensity values, 5% of transcripts (for each sample) with highest probeset variance were removed from our dataset. Applying these filters we define as cytoplasmically detected 962 filtered lncRNAs (665 genes), 906 potential protein-coding transcripts (382 genes) and 1476 protein-coding genes that are detected in K562 cytoplasm.

### RNAseq correlation analysis

ENCODE RNA-sequencing quantifications (Gencode v10 annotation) from cytoplasmic fraction of K562 cells was used to check correlation with microarray data. Correlation was calculated only with transcripts present in both ENCODE data (considered as present when RPKM bio-replicates mean different to 0 and IDR < 0.1) and microarray data.

### Classification of array transcripts

From the polysome profiling analysis, detected lncRNAs and mRNAs were classified according to the microarray sample (condition) where they displayed the highest transcript-level signal. Thus, present transcripts were classified into Heavy Polysomal (Heavy P.), Light Polysomal (Light P.) and Free Cytoplasmic transcripts (Free C.) transcripts. The remaining protein coding genes, which were not present in any microarray condition were considered not present. Remaining filtered lncRNA transcripts were subsequently checked in ENCODE K562 nucleus RNAseq. Those detected (defined as RPKM bio-replicates mean > 0 and IDR < 0.1) were classified as nuclear specific transcripts. Remaining transcripts, which are not present in cytoplasm nor in the nucleus are considered not present (NotPre).

### Cytoplasmic-nuclear localisation using RNAseq data

Cytoplasmic and nuclear RNAseq data from six different cell lines (K562, Hela, NHEK, HepG2, GM12878, HUVEC) were obtained from ENCODE (19). For each cell line we calculated cytoplasmic-nuclear RPKM ratios for transcripts detected in both that cell line and K562. RPKM was calculated as the mean of two available technical replicates, and only transcripts with mean > 0 RPKMs and IDR < 0.1 were considered present. We calculated log2 ratios of cytoplasmic expression versus nuclear expression (RPKM units) for those transcripts present in both nucleus and cytoplasm.

### Tissue Expression Analysis

We extracted tissue expression values for 16 human tissues from Human Body Map (HBM) RNAseq data, downloaded from ArrayExpress under accession number E-MTAB-513. These data were used to quantify Gencode v10 transcripts using the GRAPE pipeline(56). Transcripts were defined as ubiquitous if they had >0.1 RPKM expression in all 16 tissues.

### Transposable element analysis

The 2013 version of RepeatMasker human genomic repetitive element annotations were downloaded from UCSC Genome Browser, and was converted to BED format. Using the tool IntersectBED, we calculated (1) the number of instances of intersection, and (2) the number of nucleotides of overlap, between each lncRNA transcript and each transposable element. This analysis was carried out for both transposable element types, and transposable element classes.

### ORF analysis

We mapped all possible canonical open reading frames (ORFs) in each of six frames in lncRNA and protein coding transcripts from Gencode. If more than one start codon is in frame with a stop codon, only the start codon for the longest ORF is considered.

### CAGE analysis of lncRNA capping

5' cap analysis was performed on cap analysis gene expression (CAGE) tags from ENCODE (19) for K562 cytoplasmatic poly+ RNA and mapped these tags to the microarray region comprising between 100nt before and after transcription start sites of lncRNA. In order to assess the relationship between cytoplasmic class and capping, we compared CAGE tag presence to fractional occupancy in each class. The latter was calculated by subtracting input cytoplasmic log2 microarray expression intensity values from each of the three polysome profiling fractions intensity values (Free C., Light P. or Heavy P.). We divided transcripts into (log2) fraction occupancy bins from -2 to 2 at 0.5 bins. Transcripts with values outside this range were pooled into the last corresponding bin. Logistic regression was performed to assess the relationship between CAGE tag presence and occupancy.

### BLAST analysis of trans-homology between lncRNA and mRNA

Gencode v7 transcript-level FASTA files of mRNA (Gene type “protein coding”) and lncRNA were downloaded from Gencode. Two control sets analogous to lncRNA were also collected and processed in exactly the same way: first, Bedtools “shuffle” tool was used to extract random regions identical in size to the lncRNAs. Second, the introns of each lncRNA were concatenated, then a fragment of sequence identical in size to the mature lncRNA was extracted at a random location within this sequence. All sequences were repeat-masked using RepeatMasker with “sensitive” and “human” settings. A BLAST library was created using default settings with the mRNA sequences. LncRNA and control sequences were BLASTed against this library with maximum expectation value of 20.

### RNA Stability Assay

K562 cells were incubated with or without cyclohexamide (100ug/ml) for three hours followed by treatment with actinomycin D (5ug/ml) for 6 hours. RNA samples were taken at 0 and 6 hours following actinomycin D treatment to assess the stability of the RNA in the absence of transcription. RNA was purified using Trizol and Qiagen RNeasy columns. 1ug of RNA was used to make cDNA using RevertAid H Minus reverse transcriptase. Luminaris Color HiGreen High ROX qPCR master mix was used with gene specific primers for qRT-PCR on an ABI PRISM 7900 Sequence Detection Systems. Expression levels were normalised to the housekeeping gene GAPDH by the delta-delta Ct method.

## Acknowledgements

We thank members of the Guigo lab and the CRG Bioinformatics and Genomics Programme for many ideas and discussions, particularly Marta Melé, Joao Curado, and Marc Friedlaender, in addition to Fatima Gebauer (CRG, Gene Regulation Stem Cells and Cancer Programme). We would particularly like to thank Ferran Reverter for invaluable statistical advice. Thomas Derrien (University of Rennes) and Giovanni Bussotti (EBI) helped with evolutionary conservation analysis. We thank the CRG Genomics Core Facility, particularly Anna Ferrer, Maria Aguilar, Sarah Bonnin and Manuela Hummel, for array hybridisation and analysis. The following CRG colleagues generously helped with FISH experiments: Francois Le Dily, Maria Sanz, Carme Arnan, Maria Teresa Zomeño, as well as Timmo Zimmermann and Raquel García from CRG Microscopy Facility.

## Supplementary Data Files

**Table S1: Gencode v7 lncRNA classification.** Rows represent lncRNA transcript from Gencode v7.

Columns:

TransID: ENST ID for transcripts.

GeneID: ENSG ID for the corresponding gene.

Chr: Chromosome.

Trans_Start: Transcript start position.

Trans_End: Transcript end position.

Strand

Cellular_Localization: Classification of the transcripts into 5 different categories: 1: Present in cytoplasm (from polysome profiling experiment, K562 cell line); 2: Present in nucleus (from ENCODE nucleus RNAseq data, K562 cell line), but not in cytoplasm (polysome profiling experiment); 3: Not present in cytoplasm (polysome profiling experiment) nor in nucleus (ENCODE nucleus RNAseq data, K562 cell line); 4: Transcripts classified as potential protein coding transcripts; 5: Discarded transcripts. See Materials and Methods for details.

FreeC_intensity: Log2 intensity value for Free C. condition (NA if not present in this condition).

LightP_intensity: Log2 intensity value for light P. condition (NA if not present in this condition).

HeavyP_intensity: Log2 intensity value for Heavy P. condition (NA if not present in this condition).

WholeC_intensity:Log2 intensity value for whole cytoplasmic fraction (NA if not present in this condition).

Ribosomal_Classification: Classification for transcripts present in the cytoplasm: 1: Free C.; 2: Light P.; 3: Heavy P.

CPAT: CPAT score.

PhyloCSF: PhyloCSF score.

CPC: CPC score.

MS: information about presence of peptides in Mass Spectrometry analysis for this transcripts: 0: No peptide associated; 1: Peptide associated.

**Table S2: Correlation of gene expression quantification between microarray K562 cytoplasmic measurements, and ENCODE RNAseq data from K562 cellular fractions (19).**

**Table S3: Heavy Polysome mRNAs are most actively translated.** Shown are the numbers of ENCODE K562 mass spectrometry tags originating from ribosome-profiled mRNAs.

**Table S4: Small peptides originating from lncRNA.** Shown are the numbers of known small peptides discovered by mass spectrometry that map to Gencode v7 lncRNA (43).

**Figure S1: Comparison of lncRNA microarray and RNAseq quantifications. Steady state values for K562 cytoplasmic RNA were analysed**. RNAseq data was obtained from ENCODE. Only transcripts detected in both experiments are shown.

**Figure S2: Mean tissue expression across 16 human tissues from Human Body Map.** In the cases of potential protein coding and protein coding transcripts, data is only shown for those transcripts detected in K562.

**Figure S3: Protein interactions related to subcellular compartmentalisation.** Heatmap depicts lncRNA genes that interact with the indicated proteins, as defined by the Starbase database (57). Interactions of “Low stringency” were used in all cases. The colour scale indicates the percent difference of the actual to expected number of overlaps. The rows show lncRNA gene sets assigned to the four subcellular lncRNA categories, and the columns represent various proteins for which CLIPseq binding sites were analysed. Arrows indicate the reported subcellular localisation of the protein, identified by manual curation of the literature. We found a number of cases where the localisation of lncRNA corresponded with the known distribution of the protein to which they are bound: Nuclear-associated lncRNAs showed elevated binding to nuclear-localised proteins, including hnRNPC (P=0.007, Fisher’s exact test), U2AF65 (P=0.002), and eIFAIII (P=0.0002). In contrast, lncRNAs bound by the cytoplasmic acting IGFBP1 were significantly enriched in the Light Polysomal and free cytoplasmic fractions (P=0.007). Light Polysomal lncRNAs are enriched for binding by the TAF15 protein (P=0.033). A general depletion of Heavy Polysomal lncRNA was observed in the protein binding data.

**Figure S4: Association of ORF length with polysome density for protein coding transcripts.** Shown are histograms for % coverage of transcripts by their longest ORF. Top row: sense strand ORFs; Bottom row: antisense strand ORFs (control). Data are shown for mRNAs included in microarray design and classified by ribosomal occupancy. P values compare sense / antisense distributions in each case. Red line indicates mean ORF coverage percentage. Note the difference in mean value between Heavy and Light Polysomal means.

**Figure S5: Association of ORF length with polysome density for lncRNA transcripts.** Shown are histograms for % coverage of transcripts by their longest ORF. Top row: sense strand ORFs; Bottom row: antisense strand ORFs (control). Data are shown for all mRNAs together, for comparison. P values compare sense / antisense distributions in each case. Red line indicates mean ORF coverage percentage. Note the lack of difference in mean value between Heavy and Light Polysomal means.

**Figure S6: Gene structure characteristics of lncRNAs**. (A) Exon length distributions. (B) Intron length distributions. (C) Exon number per transcript. (D) Mature (processed) transcript length.

**Figure S7: GC content of coding and noncoding transcripts.**

**Figure S8: Polyadenylation of lncRNAs.** Using ENCODE data (19), we calculated the ratio of RPKM for PolyA+ / PolyA-K562 cytoplasmic RNAseq. RPKM values were averaged across the two available technical replicates, and only transcripts with nonzero mean values in both RNA samples retained. No statistically significant differences were found between Free Cytoplasmic lncRNAs and either of the ribosomal groups using either the Kolmogorov-Smirnov or Wilcoxon tests.

**Figure S9: Splicing efficiency of lncRNAs.** Using ENCODE data, we calculated RPKM values separately for the exons and introns of all lncRNAs. Shown are the log10 ratios of exon/intron values for all sets of transcripts. No statistically significant differences were found between Free Cytoplasmic lncRNAs and either of the ribosomal groups using either the Kolmogorov-Smirnov or Wilcoxon tests.

**Figure S10: Comparison of 5’ RNA folding energy.** Using the Vienna RNAfold programme (58) with default settings, we estimated the free energy of folding of the first 50nt of lncRNA and mRNA. While mRNA have more stable folding on average than expressed lncRNA (P=2.2e-16, Wilcoxon test), we could find no difference between either Heavy Polysomal (P=0.8) or Light Polysomal (P=0.7) and Free Cytoplasmic lncRNAs.

**Figure S11: ERVL-MaLR insertion length distributions.**

**Figure S12: The association between sense-intronic lncRNAs and nuclear localisation.** Shown is the percent of transcripts in each subcellular category that are located within the intron of a same-strand protein coding gene.

